# Using remote camera traps to assess mammal and bird assemblages across a complex environmental landscape

**DOI:** 10.1101/109538

**Authors:** Carl S. Cloyed, Laura R. Cappelli, David A. Tilson, John A. Crawford, Anthony I. Dell

## Abstract

Animals must navigate a complex mosaic of habitat types, both natural and artificial. As artificial habitats (e.g., agricultural fields) become increasingly abundant in many landscapes, species will be affected differently, depending on their habitat preferences. We investigated the diversity, richness, abundance, and biomass of mammals and birds with remote camera traps that optimized the capture of both large and small animals. Camera traps allowed us to capture natural rates of mammals and birds, which is difficult to obtain using human observers who can affect the behavior of animals and are limited in their spatio-temporal scope and ability to assess nocturnal communities. Our camera trap arrays were established along two transects in a local conservation reserve; one transect ran from an agricultural field to an upland forest and another from a wetland to an upland forest. Over the 6-week study our cameras recorded 2,245 images, within which we observed 483 individuals comprising 16 species of mammals and birds. Our data showed that species composition and abundances were only marginally different between the two transects, with species common to both transects not exhibiting any statistical difference in abundances. Coyotes and armadillos were unique to the riparian transect, and many more bird species were present along the riparian transect than the agricultural transect. Diversity, richness, and total community biomass did not differ significantly between the two transects nor along each transect but there were non-significant trends in predicted directions. Our results revealed that fewer species use the forest immediately adjacent to the agricultural field, but more species use the wetland and the forest immediately adjacent to the wetland. Our results corroborate other studies revealing that certain species are more common in forested areas but also that some species thought to prefer forested areas may actually be more habitat generalists than previously thought.

## Introduction

Animals must navigate landscapes composed of a heterogeneous mix of habitat types. As human activities modify and degrade habitats, landscapes are transforming into a mix of natural and artificial habitats [1]. More than 75% of deciduous forests in Eastern and Midwestern North America were cleared by 1850, primarily to create agricultural fields [2]. The clearing of these forests has changed the landscape matrix from primarily forested to agricultural fields [1–2]. This fragmentation of forests over the last two centuries has decreased landscape connectivity [2–4], isolated populations sensitive to these changes and increased inbreeding in those species [5–6], and produced more field-forest edges, thus increasing edge effects on forest interior species [7–9]. While these changes are detrimental to many habitat specialist species, habitat generalists often benefit from them [10]. As a result, changes in the landscape can affect species differently.

Eastern and Midwestern North American forests contain mammal and bird species that are both adversely and beneficially affected by landscape modifications. For example, white-tailed deer (*Odocoileus virginianus*) and nine-banded armadillos (*Dasypus novemcinctus*) are both forest interior species [11–14]. As such, we expect these species to be relatively infrequent in agricultural fields compared to upland and riparian forests. On the other hand, species such as raccoons (*Procyon lotor*), coyotes (*Canis latrans*), and grey squirrels (*Sciurus carolinensis*) are habitat generalists and will likely be equally common in forested and agricultural habitats [15–17]. Understanding exactly how these species respond to landscape changes and navigate a heterogeneous complex of habitat types is important for wildlife management [18].

Emerging technologies have advanced methodologies in ecology and wildlife monitoring. Automated computer imaging [19], drone technology [20–21], and motion-activated camera-traps [22–24] have provided high-throughput methods that decrease manual labor. Camera traps, for example, enable researchers to non-invasively record the abundances of a wide range of animals [25–26], including rare and endangered species [22, 27–28]. Camera traps require much less work than the manual labor required to extensively sample communities in order to properly measure species richness and abundances [23, 25], and these methods can greatly increase data throughput [19, 23, 29]. Furthermore, camera traps can provide information regarding an animal’s phenotype, including traits such as body size and body condition [29].

In this study, we used camera trap arrays to examine diversity, richness, abundance, and biomass of mammals and birds along two transects that span different ecological gradients. The first transect ran from an agricultural field to an upland forest. The second transect ran from a wetland to an upland forest. We predicted that diversity, species richness, and biomass would be higher along the riparian transect, as species that are found in both upland forests and wetlands may be present. We predicted that species compositions and abundances would be different between the transects, with forest-oriented species (white-tailed deer, armadillos) more common on the riparian transect and species that are habitat generalists (coyote, raccoons, squirrels, and Virginia opossums) found equally between the transects.

## Materials and methods

### Study site

Our study was conducted at Lindenwood University’s Daniel Boone Field Station, located in St. Charles County, MO (Lat: 38.652777; Long: -90.854376). This 404.7 hectare field station (Fig. 1) maintains an annually harvested, 6.5 hectare hay field bordered by a mixed deciduous forest dominated by black oak (*Quercus velutina*), post oak (*Quercus stellata*), and white ash (*Fraxinus americana*). Ten small wetlands are interspersed throughout the forested area, providing an important local water resource for birds and mammals during drier summer months. Climate at the field station is highly seasonal, with the average temperature during the study of 24.2 °C and daily average precipitation of 1.68 cm.

**Fig 1:**
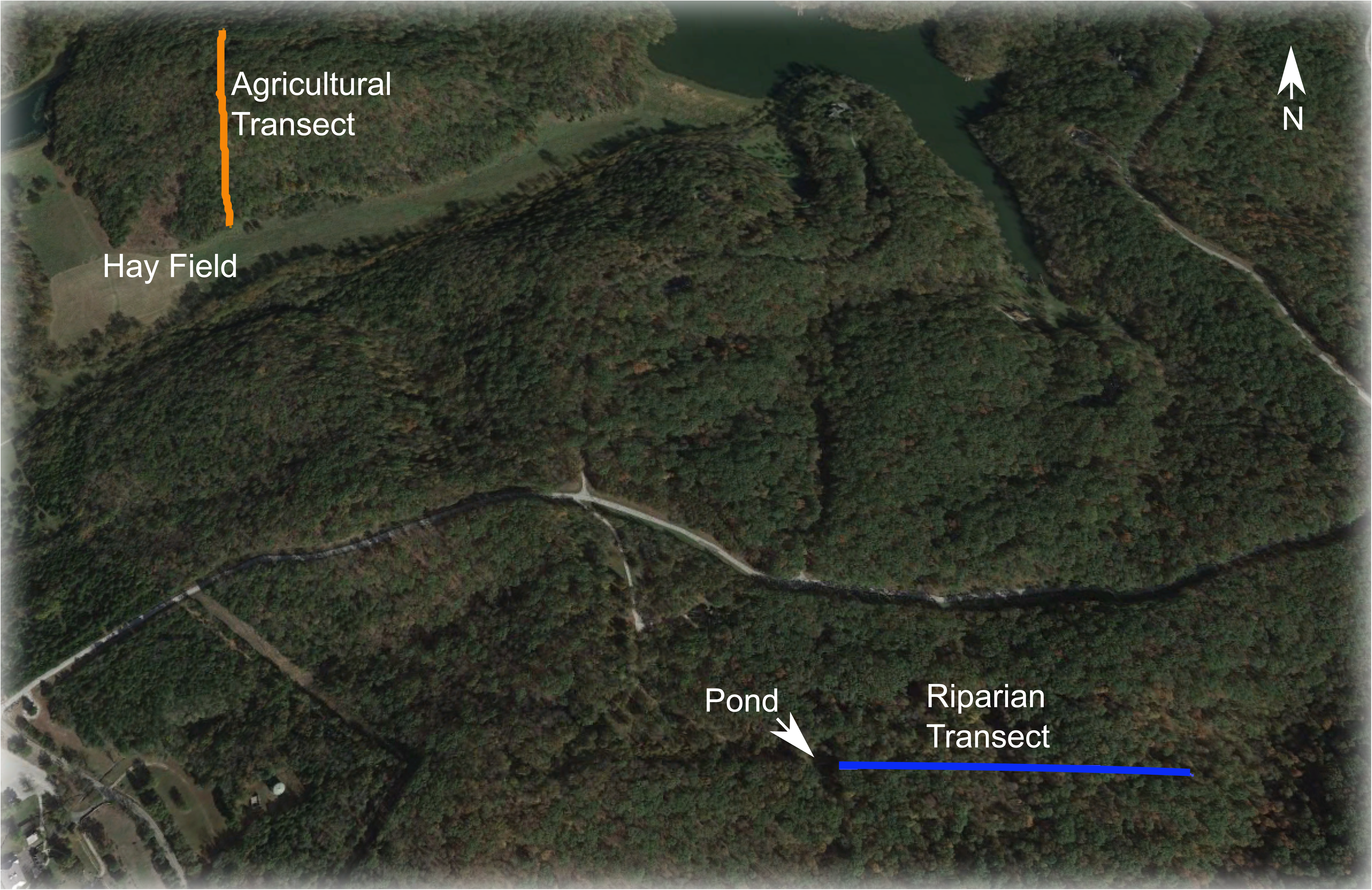
Map of the Lindenwood University’s Daniel Boone Field Station. Map shows the agricultural transect (orange) and the riparian transect (blue). The hay field is at the southern end of the agricultural transect and the small pond is at the western end of the riparian transect.

### Data collection

Remote camera traps were used to monitor mammal and bird assemblages along two 300 m transects, both positioned to span distinct environmental gradients across the field station (Fig 1). Transect 1 ran from the edge of the 6.5 hectare hay field into the surrounding upland forest, while transect 2 ran from a small wetland (~125 m^2^) into the surrounding forest. The wetland at the beginning of transect 2 was bordered by forest on three sides, and by a small, mesic meadow to the north. Puddles often formed along transect 2 after heavy rains, primarily near 142 m (Fig 1). Transect 1 and transect 2 are referred to as the agricultural and riparian transects, respectively, throughout this study.

Monitoring of each transect was conducted over a 32-day period (8 June through 10 July 2015). A total of 42 randomly-determined locations (plots) along each transect were monitored each for 48 or 72 h (depending on access to field site), with a total of three plots monitored simultaneously on each transect (the order in which plots were sampled was also randomly determined). At each plot, three infrared (IR) motion-sensor game cameras (Browning Model BTC-5) were positioned so that the fields of view overlapped, optimizing detection of small to large birds and mammals (Fig 2). Two of the cameras were attached to a single tree – one 20 cm and the other 50 cm above the substrate; both pointing in the same direction – while the third camera was placed 1–5 meters away from the first two cameras at 50 cm above the ground on a tree so that its field of view was perpendicular to the other two cameras (Fig 2). All cameras were aimed parallel to the ground, and we assume that the total area covered by the three cameras in each plot was approximately the same for all 84 plots. Vegetation in front of the cameras was removed to prevent it from obscuring the field of view. Cameras were programmed to capture images immediately after motion was detected, with subsequent photos delayed five seconds to reduce multiple images of the same individual being recorded. The field of view for each camera was 55°. The limits of the IR trigger and the IR flash illumination were 13 m and 30.5 m, respectively. Image resolution was set at 1920 x 1080 pixels, with each image also recording the time and date it was taken. This project conformed to the legal requirements for the use of vertebrates in research and was approved by the University of Illinois’ Institutional Animal Care and Use Committee (Protocol Approval # 15074).

**Fig 2:**
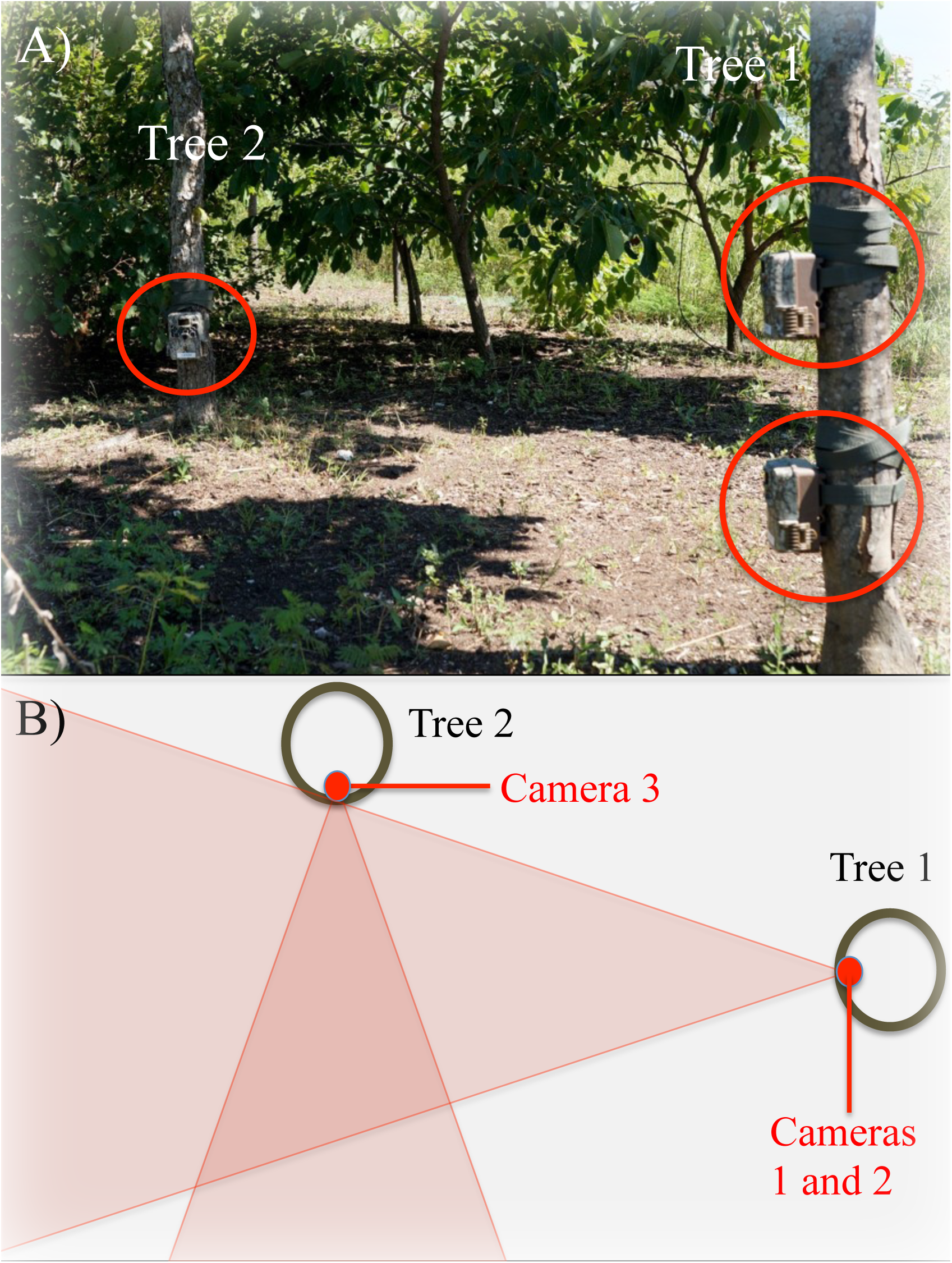
Details of our 3-camera set up at each plot. **(A)** Shows a photo of our field setup, with cameras circled in red. **(B)** Schematic of the setup from above, showing the overlap in the field of views of the three cameras at each plot. See main text for more details of our setup.

Additional environmental data was taken at each plot including: i) canopy cover (%) calculated with a spherical densiometer; ii) leaf-litter depth (mm) was the average of four random locations near the center of each plot measured with a hand ruler; and iii) environmental temperature (°C), which we measured with iButtons (Maxium Integrated, San Jose, CA) every hour. The iButtons were deployed for the duration that the camera traps were set at each plot and were used to determine the maximum and average temperatures of the plot.

### Image analysis

The taxonomic identity of all birds and mammals in each image was determined using published literature and expert opinion. While most individuals could be identified to species, squirrels (*Sciurus* spp.) and mice (*Peromyscus* spp.) were only identified to genus (for simplicity, throughout the rest of the paper we refer to these taxonomic groups as species). Images of the same species (or genus for *Sciurus* and *Peromyscus*) taken within 10 min of each other at the same plot were considered the same individual, unless this was obviously untrue (e.g., different body sizes, antlered versus non-antlered deer). The body length of all mammals was estimated from each camera trap image by comparison to standardized images from each plot that included a human observed at specific distances from the cameras. Published length-weight regressions were used to estimate body mass; for deer we used W = 0.0287L^3.03^, while for all other mammals we used W = 0.0374L^2.92^, where W = weight and L = length [30]. We did not calculate the body weight of birds because it was often difficult to determine their lengths. Mammal biomass was calculated by summing all individuals at each plot and was standardized per day. For images in which body length could not be estimated, typically because the full length of the individual was obscured, an average for that species was used (this comprised a total of 52% of individuals).

### Statistical analyses

*Environmental Variables:* Each plot was binned according to their distance from the hay field or wetland, with each bin spanning 50 m (i.e., bin 1 contained all plots between 1–50 m, bin 2 contained all plots between 51–100 m, etc., to 300 m). We then used one-way analysis of variance tests (ANOVA) to determine differences in environmental variables along each transect, where environmental variables (average temperature, maximum temperature, leaf litter depth, and percent canopy cover) were response variables, and bin number was the explanatory variable. We used two-tailed, Welch’s t-tests to determine differences in environmental variables between the two transects.

*Abundance and Composition:* To account for different sampling effort across plots, we calculated abundance as the number of individuals seen per day for each species/genus. To determine differences in species composition and abundances of each species both between and along transects we used a multi-response permutation procedure (MRPP) with a Bray-Curtis distance measure [31]. MRPP is more robust than discriminant analysis or MANOVA for species composition data, which are frequently non-normally distributed [31].

*Diversity, Richness, and Biomass:* We used general linear models (GLMs) in a model selection analysis to determine which ecological parameters best predict diversity, richness, and biomass. We used the per day abundances to estimate diversity using the Shannon-Wiener index [32]. Preliminary analysis using correlations and t-tests among the parameters revealed which factors and interaction terms were likely to be important in influencing diversity, richness, and biomass [33]. For each analysis (diversity, richness, and biomass), we began with models containing distance, transect type, maximum temperature, average temperature, litter depth, percent canopy cover, transect*distance, transect*maximum temperature, transect*litter depth, transect*percent canopy cover, distance*litter depth, and distance*percent canopy cover as explanatory variables. To determine which models best explained diversity, richness, and biomass, we used an information theoretic approach in which models were selected based on their Akaike’s information criterion corrected for small sample size (AIC_c_) [34–35]. We calculated AIC_c_ values using the *AICcmodavg* package in R [36–37]. We used the *step()* function in R to generate a subset of models [33, 37]. We considered models with the lowest AIC_c_ values to be the models of best fit, and we ranked models based on ΔAIC_c_, where ΔAIC_c_ = AIC_i_ – AIC_m_ and AIC_i_ is the AIC_c_ value of model *i* and AIC_m_ is the AIC_c_ value of the model of best fit. When the ΔAIC_c_ was less than two, indicating that both models substantially supported the data, we used the normalized Akaike model weights (*w_im_*) to determine the best model, where 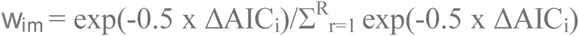 [35] The normalized weight of each of the best models was less than 0.9, indicating other models were supported by the data, so we performed model averaging which provides more robust model variances and increases the reliability of parameter estimates [34]. We included a final subset of models that had cumulative model weights of ≥ 0.95 [38]. To determine the relative importance of each term in the models, we calculated the normalized Akaike weight for each parameter (*w_ip_*), which is the sum of the *w_im_* in which that parameter is present (*w_ip_* = 1 indicates a parameter present in all models). We calculated the confidence intervals of the slopes between each parameter and diversity, species richness, and biomass to determine when those parameters may have significant effects. Additionally, we performed ANOVAs to determine if diversity, richness, and biomass differed along each transect (see above section “*environmental variables*” for binning procedure), and we used two-sided, Welch’s t-tests to determine differences between transects.

## Results

### Environmental variables

Average temperature was higher on the agricultural transect than on the riparian transect (t-test; t = 2.149, df = 74.69, *p* = 0.035). Average temperature varied along the agricultural transect (Fig 3A; ANOVA: F = 3.124, df = 5, 32, *p* = 0.021), with significant differences occurring between bin two (51–100 m) and bin six (251–300 m; *p* = 0.027) and between bin two and bin three (101–150 m; *p* = 0.030). Average temperatures did not differ along the riparian transect (Fig 3A; F = 0.282, df = 1, 37, *p* = 0.598). Maximum temperature was higher on the agricultural transect (Figure 3B; t-test: t = 2.893, df = 70.41, *p* = 0.005). Maximum temperature did not vary with distance along the agricultural (Fig 3B; F = 1.233, df = 5, 32, *p* = 0.317) or the riparian transect (Figure 3B; ANOVA: F = 1.218, df = 1, 37, *p* = 0.277). Litter depth did not vary between the transects (Fig 3C; t-test: t = 0.428, df = 74.99, *p* = 0.670). Litter depth did increase with distance from the hay field (Fig 3C; ANOVA: F = 4.533, df = 5, 32, *p* = 0.003). Along the agricultural transect, a Tukey’s Honest Significant Differences test found that litter depths were significantly different between bin one (1–50 m) and bin four (151–200 m; *p* = 0.053), bin one and bin five (201–250 m; *p* = 0.007), and bin one and bin six (251–300 m; *p* = 0.003). There was no change in litter depth along the riparian transect (Fig 3C; ANOVA: F = 2.554, df = 1, 37, *p* = 0.119). The canopy cover was greater along the riparian transect (Fig 3D; t-test: t = –2.397, df = 68.08; *p* = 0.019). Along the agricultural transect, canopy cover was lowest nearest the hay field (Fig 3D; ANOVA: F = 4.443, df = 5, 32, *p* = 0.003). The canopy cover was significantly different between bin one and every other bin number (bin two: *p* = 0.021; bin three: *p* = 0.002; bin four: *p* = 0.013; bin five: *p* = 0.038; bin six: *p* = 0.007). The canopy cover did not change along the riparian transect (Fig 3D; ANOVA: F = 0.446, df = 1, 37, *p* = 0.508).

**Fig 3:**
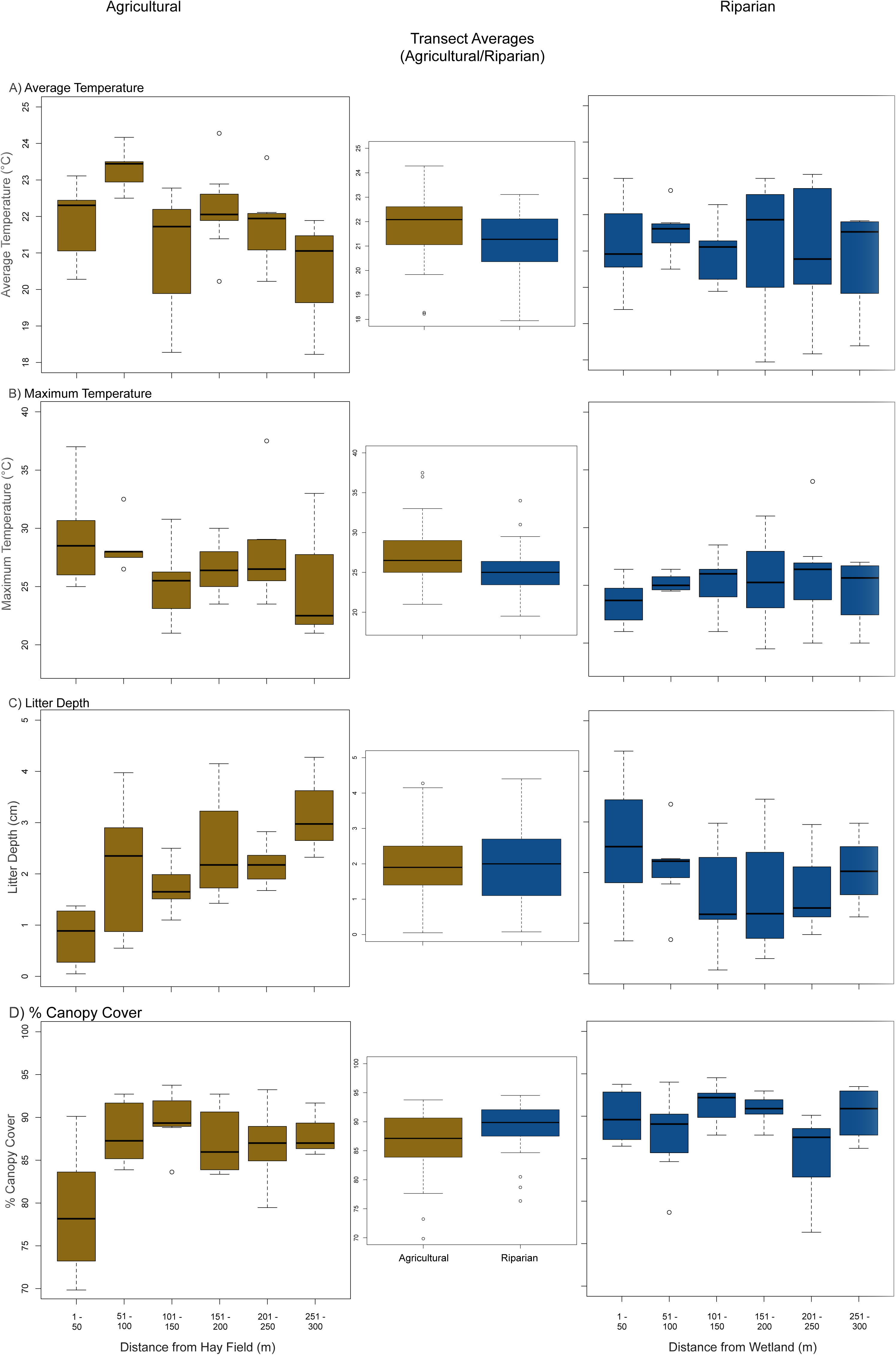
Environmental variables along and between transects. Box plots of environmental variables at different distances along the agricultural (left side) and riparian (right side) transect, with comparisons between the two transects in the middle.

### Species Composition and Abundances

We recorded at least one mammal or bird from 92 % of the plots we sampled; 38 of 42 plots along the agricultural transect and 39 of 42 plots along the riparian transect. In total, we captured 2,245 images that included at least one mammal or bird, which we estimated to represent 483 individuals. For mammals, this consisted of 231 squirrels (*Sciurus* spp.), 81 white-tailed deer (*Odocoileus virginianus*), 64 Virginia opossums (*Didelphis virginiana*), 36 deer mice (*Peromyscus* spp.), 38 raccoons (*Procyon lotor*), 6 armadillos (*Dasypus novemcinctus*), and 3 coyotes (*Canis latrans*). There were a total of 24 birds captured, with tufted titmouse (*Baeolophus bicolor*), wild turkeys (*Meleagris gallopavo*), and the northern cardinal (*Cardinalis cardinalis*), occurring on both transects and coopers hawks (*Accipiter cooperii*), blue jays (*Cyanocitta cristata*), wood thrushes (*Hylocichla mustelina),* and barred owls (*Strix varia*) found only on the riparian transect. Due to their low numbers, all bird species were included as a single taxonomic group in subsequent analyses. Using a MRPP, we found a marginally significant difference in species composition and abundances between the two transects (A = 0.0077, *p* = 0.0706). Squirrels (the most common species on both transects), white-tailed deer and raccoons exhibited similar per day abundances between our two transects (Table 1; Fig 4A, B). Virginia opossums and mice were both twice as abundant on the agricultural transect, whereas birds were over three times more abundant on the riparian transect (Table 1; Fig 4A, B). Armadillos and coyotes were recorded only on the riparian transect (Table 1; Fig 4A, B), and no species were found exclusively on the agricultural transect.

**Fig 4:**
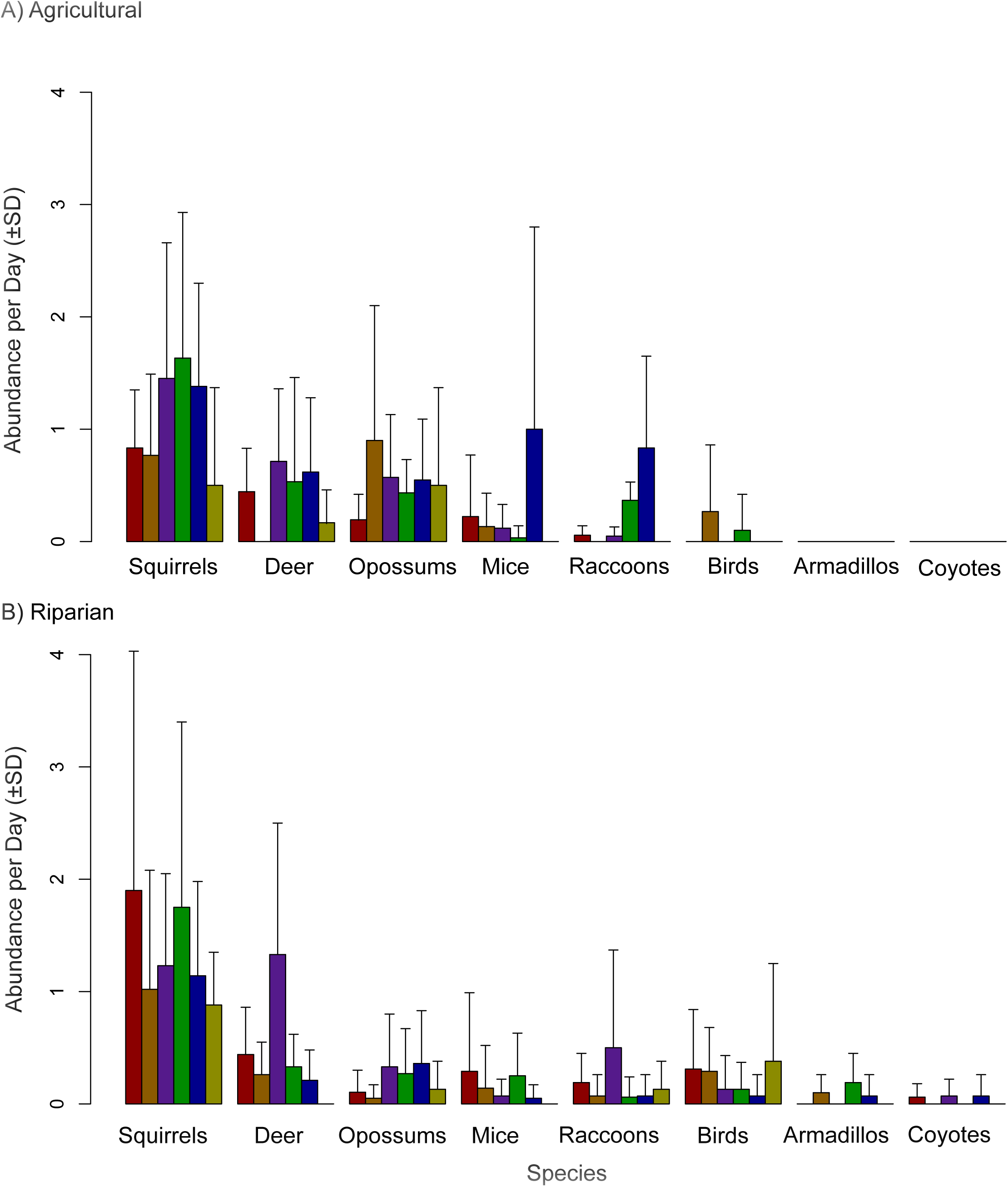
Species abundances on along each transect. Abundance per day (±SD) of each species/genus at different distances along the A) agricultural and B) Riparian transects. Each color represents a different distance along the transects. Red = 1-50m, orange = 51-100m, purple = 101-150m, green = 151-200m, blue = 201-250m, and yellow = 251-300m.

**Table 1.**
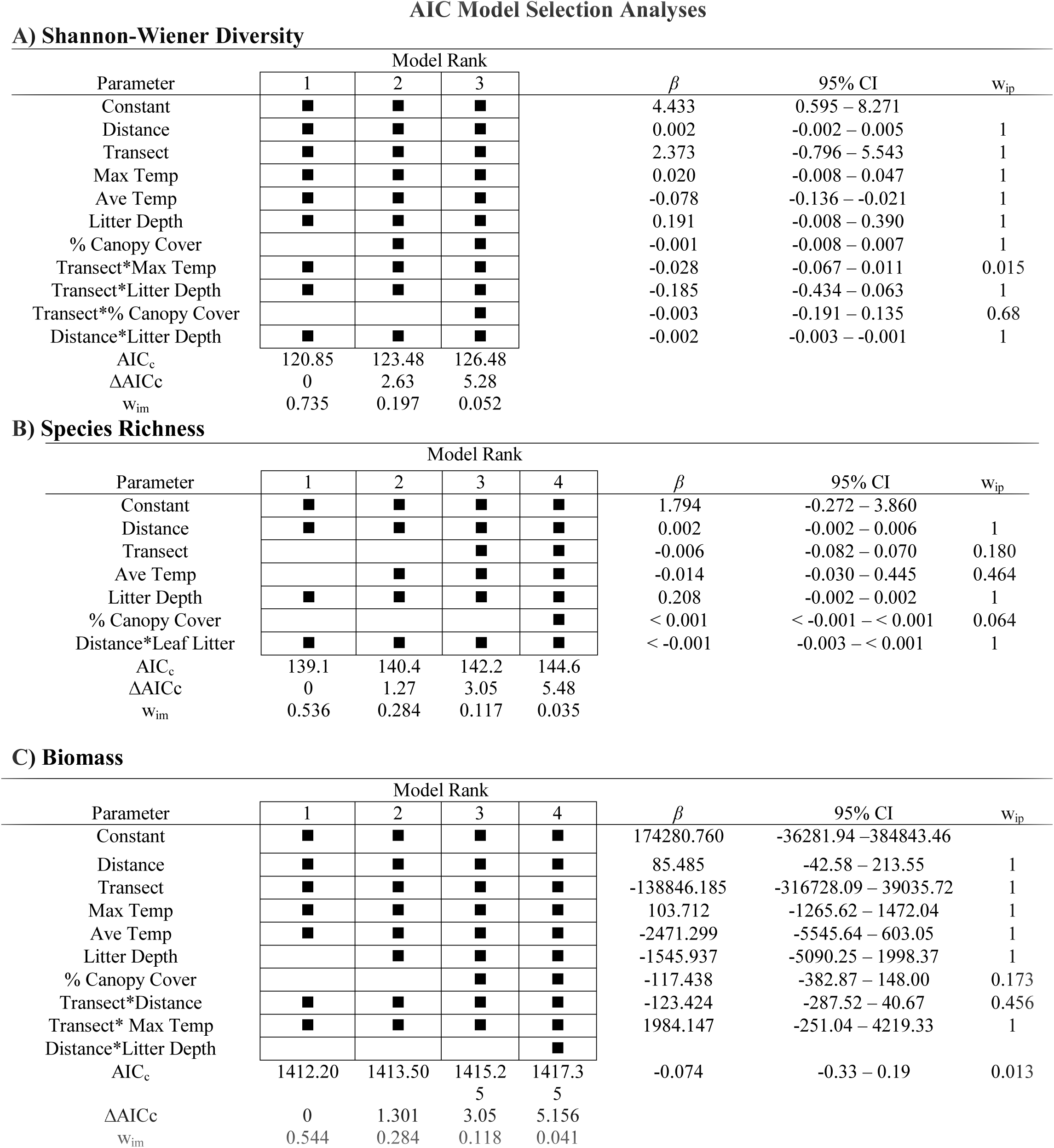
AIC_c_ model selection analysis for A) Shannon-Wiener diversity indices, B) Species Richness, and Biomass for the subset of models that have a cumulative model weight of 0.95 and model-averaged parame analysis. The black symbols indicate the presence of that parameter in the model. The AIC_c_ values, ΔAIC_c_, model weights (w_im_) are found on the lower part of the table. We used model-averaging to obtain parameter estimates, which are displayed on the right section of the table.

A) Shannon-Wiener Diversity
| Parameter | Model Rank | | | $\beta$ | 95% CI | $w_{ip}$ |
| --- | --- | --- | --- | --- | --- | --- |
|  | 1 | 2 | 3 |  |  |  |
| Constant | ■ | ■ | ■ | 4.433 | 0.595 – 8.271 |  |
| Distance | ■ | ■ | ■ | 0.002 | -0.002 – 0.005 | 1 |
| Transect | ■ | ■ | ■ | 2.373 | -0.796 – 5.543 | 1 |
| Max Temp | ■ | ■ | ■ | 0.020 | -0.008 – 0.047 | 1 |
| Ave Temp | ■ | ■ | ■ | -0.078 | -0.136 – -0.021 | 1 |
| Litter Depth | ■ | ■ | ■ | 0.191 | -0.008 – 0.390 | 1 |
| % Canopy Cover |  | ■ | ■ | -0.001 | -0.008 – 0.007 | 1 |
| Transect*Max Temp | ■ | ■ | ■ | -0.028 | -0.067 – 0.011 | 0.015 |
| Transect*Litter Depth | ■ | ■ | ■ | -0.185 | -0.434 – 0.063 | 1 |
| Transect*% Canopy Cover |  |  | ■ | -0.003 | -0.191 – 0.135 | 0.68 |
| Distance*Litter Depth | ■ | ■ | ■ | -0.002 | -0.003 – -0.001 | 1 |
| AIC<sub>c</sub> | 120.85 | 123.48 | 126.48 |  |  |  |
| ΔAIC<sub>c</sub> | 0 | 2.63 | 5.28 | | | |
| $w_{im}$ | 0.735 | 0.197 | 0.052 | | | |

B) Species Richness
| Parameter | Model Rank | | | | $\beta$ | 95% CI | $w_{ip}$ |
| --- | --- | --- | --- | --- | --- | --- | --- |
|  | 1 | 2 | 3 | 4 |  |  |  |
| Constant | ■ | ■ | ■ | ■ | 1.794 | -0.272 – 3.860 |  |
| Distance | ■ | ■ | ■ | ■ | 0.002 | -0.002 – 0.006 | 1 |
| Transect |  |  | ■ | ■ | -0.006 | -0.082 – 0.070 | 0.180 |
| Ave Temp |  | ■ | ■ | ■ | -0.014 | -0.030 – 0.445 | 0.464 |
| Litter Depth | ■ | ■ | ■ | ■ | 0.208 | -0.002 – 0.002 | 1 |
| % Canopy Cover |  |  |  | ■ | < 0.001 | < -0.001 – < 0.001 | 0.064 |
| Distance*Leaf Litter | ■ | ■ | ■ | ■ | < -0.001 | -0.003 – < 0.001 | 1 |
| AIC<sub>c</sub> | 139.1 | 140.4 | 142.2 | 144.6 |  |  |  |
| ΔAIC<sub>c</sub> | 0 | 1.27 | 3.05 | 5.48 | | | |
| $w_{im}$ | 0.536 | 0.284 | 0.117 | 0.035 | | | |

**Table 1.** AIC<sub>c</sub> model selection analysis for A) Shannon-Wiener diversity indices, B) Species Richness, and Biomass for the subset of models that have a cumulative model weight of 0.95 and model-averaged parameter analysis.
| Parameter | Model Rank | | | | $\beta$ | 95% CI | $w_{ip}$ |
| --- | --- | --- | --- | --- | --- | --- | --- |
|  | 1 | 2 | 3 | 4 |  |  |  |
| Constant | ■ | ■ | ■ | ■ | 174280.760 | -36281.94 – 384843.46 |  |
| Distance | ■ | ■ | ■ | ■ | 85.485 | -42.58 – 213.55 | 1 |
| Transect | ■ | ■ | ■ | ■ | -138846.185 | -316728.09 – 39035.72 | 1 |
| Max Temp | ■ | ■ | ■ | ■ | 103.712 | -1265.62 – 1472.04 | 1 |
| Ave Temp | ■ | ■ | ■ | ■ | -2471.299 | -5545.64 – 603.05 | 1 |
| Litter Depth |  | ■ | ■ | ■ | -1545.937 | -5090.25 – 1998.37 | 1 |
| % Canopy Cover |  |  | ■ | ■ | -117.438 | -382.87 – 148.00 | 0.173 |
| Transect*Distance | ■ | ■ | ■ | ■ | -123.424 | -287.52 – 40.67 | 0.456 |
| Transect* Max Temp | ■ | ■ | ■ | ■ | 1984.147 | -251.04 – 4219.33 | 1 |
| Distance*Litter Depth |  |  |  | ■ |  |  |  |
| AIC<sub>c</sub> | 1412.20 | 1413.50 | 1415.2 | 1417.3 | -0.074 | -0.33 – 0.19 | 0.013 |
|  |  |  | 5 | 5 |  |  |  |
| ΔAIC<sub>c</sub> | 0 | 1.301 | 3.05 | 5.156 | | | |
| $w_{im}$ | 0.544 | 0.284 | 0.118 | 0.041 | | | |

Along the agricultural transect, squirrels, white-tailed deer, raccoons, and birds were most abundant at intermediate distances from the hay field (Fig 4A). Virginia opossums were uncommon nearest the hay field edge, becoming more abundant moving away from the hay field (Fig 4A). Mice were most common farther from the hay field, although they also had relatively high abundances near the hay field (Fig 4A). Along the riparian transect, white-tailed deer, Virginia opossums, raccoons, and armadillos had the highest abundances at intermediate distances from the wetland (Fig 4B). Squirrels and mice had two distinct modes of abundance, near the wetland and at intermediate distances from the wetland (Fig 4B). Birds were most abundant near the wetland and at plots farthest from the wetland (Fig 4B). However, despite apparent systematic differences in the abundances of different species/genera at different distances along each transect, a MRPP found no difference in species composition and abundances at different distances along either the agricultural transect (A = –0.0079, *p* = 0.6065) or the riparian transect (A = 0.0021, *p* = 0.6894).

### Diversity, species richness and biomass

The model selection process varied between diversity, species richness, and biomass (Table 1A, B, C, respectively). In the model selection for diversity, three models had cumulative weights ≥ 0.95 (Table 1A), with the best-fitting model containing distance, transect, maximum temperature, average temperature, litter depth, transect*maximum temperature, transect*litter depth, and distance*litter depth. The model weight was less than 0.9 for this model, but the next best-fitting model, which included canopy cover, had much lower weight and a ΔAIC_c_ above two (Table 1A). In the model selection analysis for species richness, four models had cumulative weights of ≥ 0.95 (Table 1B), with the best-fitting model containing distance, litter depth, and the distance-litter depth interaction. The best-fitting model had a comparably low model weight than the second best-fitting model, which included average temperature, and the ΔAIC_c_ value of the second best-fitting model was less than two. The model selection for biomass produced four models with cumulative model weights ≥ 0.95 (Table 1C), with the best-fitting model containing distance, transect, maximum temperature, average temperature, transect*distance, transect*maximum temperature (Table 1C). The second best-fitting model, which included litter depth, had a comparably high model weight and a ΔAIC_c_ value less than two. Distance is the only parameter common to the best fitting models for diversity, richness, and biomass. Common parameters between diversity and richness include distance, litter depth, and the distance-litter depth interaction.

The model selection analysis for diversity included eight explanatory parameters in the model of best fit (Table 1A). Diversity decreased at plots with higher average temperatures along both the agricultural and riparian transects (Fig 5A), but only decreased along the riparian transect for maximum temperature (Fig 5B). Diversity also decreased with increasing litter depth along both transects (Fig 5C). Despite being included in the best-fitting model, the slopes of these variables were not significantly different from zero (Table 1A). Likewise, diversity did not differ significantly between the agricultural and riparian transects (Fig 6A; t-test: t = 0.214, df = 74.08, *p* = 0.831). Diversity did differ by bin number along the agricultural transects (Fig 6A; ANOVA: F = 2.742, df = 5, 32, *p* 0.036). Follow-up Tukey Honest Significant Difference tests found differences in diversity between bins three and six (101–151 m and 251–300 m from the hay field; *p* = 0.048). Diversity did not differ among bin numbers on the riparian transect (Fig 6A: F = 0.278, df = 5, 32, *p* = 0.922).

**Fig 5:**
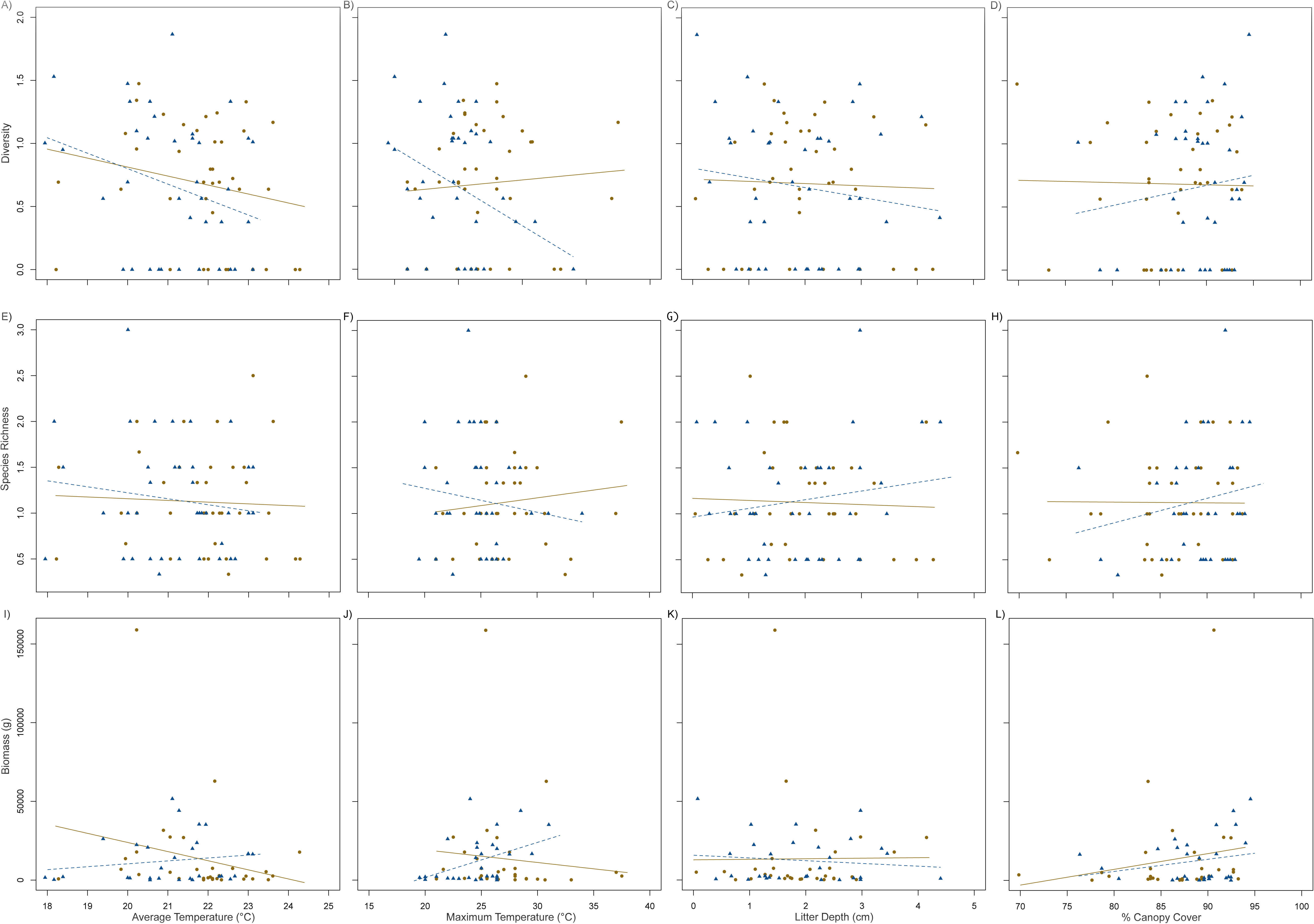
Effects of environmental variables on diversity, species richness, and biomass. The effects of average temperature, maximum temperature, litter depth, and percent canopy cover on diversity (**A-D**) and species richness (**E-H**), and biomass (**I-L)** of birds and mammals. Solid circles denote individuals observed on the agricultural transect, and open triangles denote individuals observed on the riparian transect. Lines represent the best-fit GLM for agricultural (solid) and riparian (dashed) transects.

**Figure 6:**
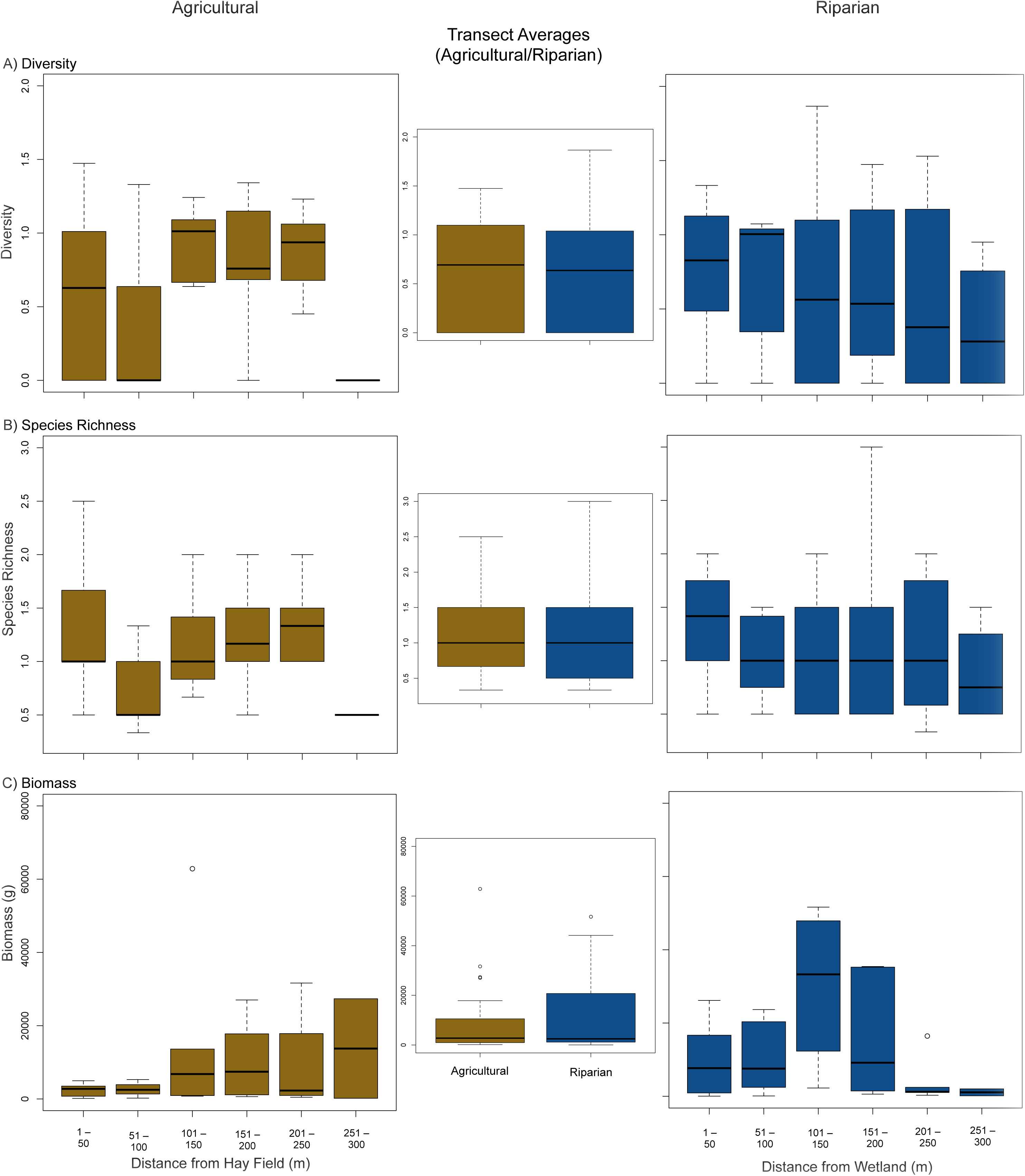
Diversity, species richness, and biomass along and between each transect. A) Diversity, B) species richness, and C) Biomass along the agricultural and riparian transects (left and right columns, respectively) and between the agricultural and riparian transects (middle column).

The model selection analysis for species richness included three parameters for the model of best fit (Table 1B). Species richness increased slightly with increased litter depth along the riparian transect but not the agricultural transect (Figure 5G). The difference in these slopes was not significant though (Table 1B). Species richness did not differ between the agricultural and riparian transects (Fig 6B; t-test: t = –0.171, df = 73.93, *p* = 0.864). Species richness did not differ with bin number along either the agricultural (Figure 6B; ANOVA: F = 2.017, df = 5, 32, *p* = 0.103) or riparian transects (Fig 6B; ANOVA: F = 0.352, df 5, 32, *p* = 0.877).

The model selection analysis for biomass included six parameters in the model of best fit (Table 1C). Biomass decreased at plots with maximum temperatures along the agricultural transect but increased with maximum temperature on the riparian transect (Fig 6C). The difference in these slopes was not significant however (Table 1C). Biomass did not differ between the agricultural and riparian transects (Fig 6C; t-test: t = 0.182, df = 43.77, *p* = 0.856) did not differ with bin number along either the agricultural (Fig 6C; ANOVA: F = 0.447, df = 5, 25, *p* = 0.811) or the riparian transects (Fig 6C; ANOVA: F = 2.173, df = 5, 24, *p* = 0.091).

## Discussion

Using remote camera traps, we estimated the abundance, diversity, and biomass of mammals and birds along two transects that traversed two distinct environmental gradients. Our use of multiple cameras per plot – positioned at different heights above the forest floor – increased our ability to record both small and large species. Cameras placed at standard heights above the ground (~50 cm) often do not capture small mammals and birds [39,40], while cameras placed lower can miss larger mammals [40]. Having multiple cameras with overlapping fields of view also reduced the probability of missing animals due to failure of a single camera to trigger, which occurs for a variety of reasons [27, 41]. Thus, our multiple-camera setup (Fig 2) allowed us to estimate abundance, diversity, and biomass across a wider range of mammal and bird species than would be possible with just a single camera at each plot.

There were no differences in the abundances of species that were found on both transects. The marginally significant difference in species composition and abundance found by the MRPP between the transects was likely driven by the presence of armadillos and coyotes on the riparian transect but not on the agricultural transect as well as by the increased number of birds on the riparian transect. Several of our predictions regarding species-specific trends in abundances were supported, despite no significant differences in the abundances of most mammal species between the two transects (Fig 4A, B). Armadillos were only present on the riparian transect, corroborating studies that found armadillos avoid open habitats and grasslands [11, 14]. The two species of squirrels at our site are considered habitat generalists and are quite adaptable to novel environments [15, 42–45] so it was unsurprising they occurred in high abundance on both transects (Table 1). Similarly, raccoons are a common habitat generalist in the Midwestern United States [17] and were found nearly equally between the transects.

Several of our other predictions regarding abundances were not supported. White-tailed deer, a species often associated with forests [12–13], were captured about equally on the two transects. While white-tailed deer prefer forested habitats, they will still use a wide range of habitats, including open fields [45]. Our study site is outside the range of mule deer, *Odocoileus hemionus*, which are closely related to white-tailed deer and found in more open areas in the western half of North America. Without the presence of mule deer, white-tailed deer may utilize open habitats and forests adjacent to open habitats more frequently than when in the presence of mule deer. Coyotes were found only on the riparian transect in our study. Coyotes are habitat generalists that will use habitats associated with their prey [16–17]. Coyotes have been shown to be one of the few species that did not avoid agricultural fields [8], and it was therefore surprising that they were not captured on the agricultural transect, where their prey were equally abundant as on the riparian transect. However, coyotes may avoid cameras [39], making it difficult to conclude any patterns in their habitat use. Even on the riparian transect, coyotes were captured only three times. Virginia opossums tended to be more abundant along the agricultural transect, contrary to our prediction that they would be found about equally between the transects (Fig 4A, B).

Contrary to our prediction and despite higher species richness on the riparian transect, diversity was nearly equal between the two transects (Fig 6A). Two main reasons explain this pattern of diversity. First, the distribution of species is more even along the agricultural transect (Fig 4A, B). Squirrels dominate in abundance on both transects, but are relatively more dominant on the riparian transect than on the agricultural transect (Fig 4A, B). This unevenness in species abundances likely overrides the presence of two more species on the riparian transect. Second, two species of squirrels and many species of birds were respectively lumped into squirrel or bird groups in the diversity analysis. Lumping birds into a single group results in a greater underestimation in diversity on the riparian transect than on the agricultural transect. On the agricultural transect, the only birds captured were the tufted titmouse, wild turkeys, and the northern cardinal. In addition to those species, several coopers hawks, blue jays, wood thrushes, and barred owls were found on the riparian transect.

The parameters in the best-fitting model for diversity included distance, transect, maximum and average temperature, leaf-litter depth, transect*maximum temperature, transect*leaf-litter depth, and distance*leaf-litter depth. Of those parameters, average temperature and the distance*leaf-litter depth had the strongest effects, in which the 95% CI of the slopes for each parameter did not cross zero (Table 1A). Diversity decreased with increased average and maximum temperatures (Fig 5A, B), a trend that is often found with increasing temperatures [46–47]. Fewer species may be able to tolerate warmer temperatures and therefore warmer plots had fewer species. Diversity also decreased with leaf-litter depth, particularly on the riparian transect (Fig 5C). This may be driven by squirrel abundances at sites with more leaf litter [44]. Where squirrel abundances are highest, they may dominate in the diversity index and result in lower diversity at those sites. Mice may be less abundant where leaf litter is deep because they avoid dry hardwood litter to reduce the risk of auditory detection by predators [49].

The parameters in the best-fitting model for biomass included distance, transect, maximum and average temperature, transect*distance, and transect*maximum temperature. The slopes of all these parameters had 95% CI that included zero, indicating that none of the parameters significantly affected biomass. However, biomass increased gradually from the hay field to the upland forest (Fig 6E), a pattern we would expect if fewer species are using the hay field [8]. The diversity of mammals and birds found near the hay field was relatively low (Fig 6B). The concentration of biomass farther from the hay field and the lower species diversity supports the idea that the field may not be used by many species. On the other hand, biomass on the riparian transect was higher near the wetland and at intermediate distances (Fig 6F), and the diversity on the riparian transect decreased with distance from the wetland (Fig 6C), supporting the idea that many species are using the wetlands or areas adjacent to the wetlands. Wetlands are likely more useful for many organisms than the hay field. The wetland provides water, potentially a greater diversity of plants for herbivores to consume, and a greater diversity and abundance of prey for predator species. In fact, only on the riparian transect did we observe carnivores.

The relationship between biomass and temperature was opposite between the transects (Fig 5E, F). Biomass decreased with temperature on the agricultural transects and increased with temperature on the riparian transect. This pattern was more pronounced for maximum temperature (Table 3; Fig 5E, F), which varied more between the transects than average temperature (Fig 3D, E). Biomass was likely lower at warmer temperatures along the agricultural transects because plots with higher maximum temperatures were found near the hay field, where biomass was low.

An important caveat in the use of camera traps for wildlife studies is that it can be difficult to accurately determine density and abundances of animals that cannot be individually identified. Species with unique individual markings (e.g., tigers and jaguars) can be used in mark-recapture methods [22, 49]. When researchers are unable to identify individuals from markings, it becomes impossible to determine if each photograph represents separate individuals. However, researchers can take measures to reduce counting an individual more than once. We set our camera traps to have a five seconddelay after a photo was taken and when more than one image of the same species (or genus for *Sciurus* and *Peromyscus*) were taken within 10 min of each other at the same plot, they were considered the same individual, unless they were was obviously different individuals (e.g., different body size, antlered versus non-antlered deer). Many researchers have compared camera-trapping methods with traditional methods and have found that camera trapping provides comparable and reliable results [23–24, 50]. Others have found that camera traps under-estimated abundances and richness compared to traditional methods but these under-estimations were not significantly different from traditional methods [28, 51]. When researchers are only interested in detecting species presence and determining species richness, camera trapping is often the better method [25–26]. As such, the risk of resampling the same individuals more than once does not appear to affect the results any more than it does with traditional methods.

Camera traps are proving to be an invaluable tool in the toolbox of ecologists and wildlife biologists. They represent a cost-effective method for determining abundances, diversity, and richness [25, 52]. Camera traps provide the best results when they are applied to animals that can be individual identified, but still provide comparably reliable estimates on “unmarked” individuals. Indeed, their ability to capture rare and cryptic species far out-stripes traditional methods. Finally, while camera traps have been less utilized to non-invasively determine phenotypic traits, their ability to do so can greatly increase their capabilities as an ecological tool [29]. In our study, we used camera traps to estimate body size and, hence, the amount of biomass moving through each of the plots. As far as we know, this is the first study to utilize camera traps to obtain body sizes, and we conclude that future studies using camera traps can obtain body size and incorporate that information into their analyses.

In conclusion, we found that camera traps worked to estimate diversity, richness, abundance, and biomass of large and small mammals and birds. We found slight, non-significant differences in diversity, abundances, and biomass between a transect that traverses from a wetland to an upland forest and a transect that traverses from an agricultural field to an upland forest in eastern Missouri. Fewer species were found near the hay field while more species were found near the wetland. Armadillos and coyotes were unique to the riparian transect, while birds were more common and diverse on the riparian transect. Virginia opossums were more common along the agricultural transects, but no species were unique to the agricultural transect.

## Author Contributions

CSC contributed to the investigation, methodology, supervision, formal analysis, visualization, validation, and writing of the manuscript (both the original draft and review/editing).

LRC contributed to the investigation and methodology.

DAT contributed to the investigation and methodology.

JAC contributed to funding acquisition, methodology, resources, validation, and reviewing/editing the manuscript.

AID contributed to funding acquisition, methodology, resources, validation, and reviewing/editing the manuscript.

## Acknowledgments

We thank Lindenwood University for access to the field sites.

